# EEG correlates of olfactory processing during an instructed-delay task

**DOI:** 10.1101/2022.12.06.519284

**Authors:** Ivan Ninenko, Daria F. Kleeva, Nikita Bukreev, Mikhail A. Lebedev

## Abstract

EEG correlates of olfaction are of fundamental and practical interest for many reasons. In the field of neural technologies, olfactory-based brain-computer interfaces (BCIs) represent an approach that could be useful for neurorehabilitation of anosmia, dysosmia and hyposmia. While the idea of a BCI that decodes neural responses to different odors and/or enables odor-based neurofeedback is appealing, the results of previous EEG investigations into the olfactory domain are rather inconsistent, particularly when non-primary processing of olfactory signals is concerned. Here we report the results from an EEG study where an olfaction-based instructed-delay task was implemented. We utilized an olfactory display and a sensor of respiration to present odors in a strictly controlled fashion. Spatio-spectral EEG properties were analyzed to assess the olfactory-related components. Under these experimental conditions, EEG components representing processing of olfactory information were found around channel C4, where changes were detected for the 12-Hz spectral component. Additionally, significant differences in EEG patterns were observed when responses to odors were compared to the responses to odorless stimuli. We conclude EEG recordings are suitable for detecting active processing of odors. As such they could be integrated in a BCI that strives to rehabilitate olfactory disabilities or uses odors for hedonistic purposes.

## 1 Introduction

Electroencephalography (EEG) is a well-developed method for non-invasive research of cortical processing. While the EEG approach has been successfully used in the studies of motor control, visual and auditory processing, the results of EEG investigations into the olfactory domain are rather mixed. In a pioneering study, Moncrief (1962) reported that a wide range of odors (essential oils, synthetic oils and unpleasant chemicals) produced a general reduction in alpha activity, except for ylang-ylang that did not produce any effect. This observation was not fully supported by further studies, with some researchers reporting an increase in alpha activity (van Toller et al,1993) and others reported no significant change. Increases in theta activity were also reported (Klemm et al, 1993). A review of 13 papers in this field (Martin, 1998) concluded that the currently available literature paints a rather sketchy and inconclusive picture of how olfactory processing could be assessed with EEG recordings. These limitations of the application of EEG methodology to olfaction impede the development of olfactory-based BCIs that potentially could be very useful to treat olfactory disabilities, such as anosmia, dysosmia and hyposmia.

Despite the absence of agreement on the effects of odors upon EEG, several research groups reported successful attempts to use neural networks to classify perceived odors based on the recorded EEG data (Aydemir, 2017; Abbasi et al., 2019; Ezzatdoost, Hojjati and Aghajan, 2020; Saha et al 2013). All these studies relied on manual delivery of odors to participants. In this approach, odorants are usually stored in unlabelled bottles or on scented paper and then manually brought to the participant’s nose by the experimenter. Such manual delivery of odorants has many limitations, such as experimenter bias and uncontrolled amount of odorant (Lorig, 2000). Additionally, hand movement in direct proximity of the subject’s face can affect participants’ perception and add additional confounds to the experimental paradigm.

Several groups explored chemosensory event-related potentials (CSERPs) and olfactory event-related potentials (OERPs) to gain knowledge of how olfactory information is processed in the brain (Murphy et al, 2000; Rombaux et al, 2006; Invitto, Mazzatenta, 2019). In these studies, precise timing of odor onset was required to study event-related potentials (ERPs). For this purpose, a special tool was used called olfactometer. An olfactometer consists of a tube and/or a facial mask that releases odorants close to the nose.

Here we employed a new experimental paradigm for olfactory perception in human participants, which solves the problems described above and paves way toward an olfactory-based BCI. We developed an experimental paradigm where olfactory stimuli were incorporated in an instructed-delay task and electrophysiological recordings were conducted based on the most recent methodological recommendations for EEG studies of olfaction (Lorig, 2000). Automated odor delivery system was implemented that enacted near-natural olfactory experiences. We collected EEG and respiration data. An analysis of spatio-spectral EEG properties revealed neural patterns exhibited during this instructed-delay odor-discrimination task.

## 2 Materials and Equipment

### 2.1 Controlled delivery of odors

For controlled and near-natural presentation of olfactory stimuli, we utilized an automated odor delivery system developed by Sensorylab Inc. The system used piezoelectric transducers to release liquid odorants into the stream of air. (Fig. 1)The device controlled the amount of odorant by regulating stimulus onset and offset. In this system, an odorant reached the subject approximately 400 ms after stimulus onset.

**Figure 1.**
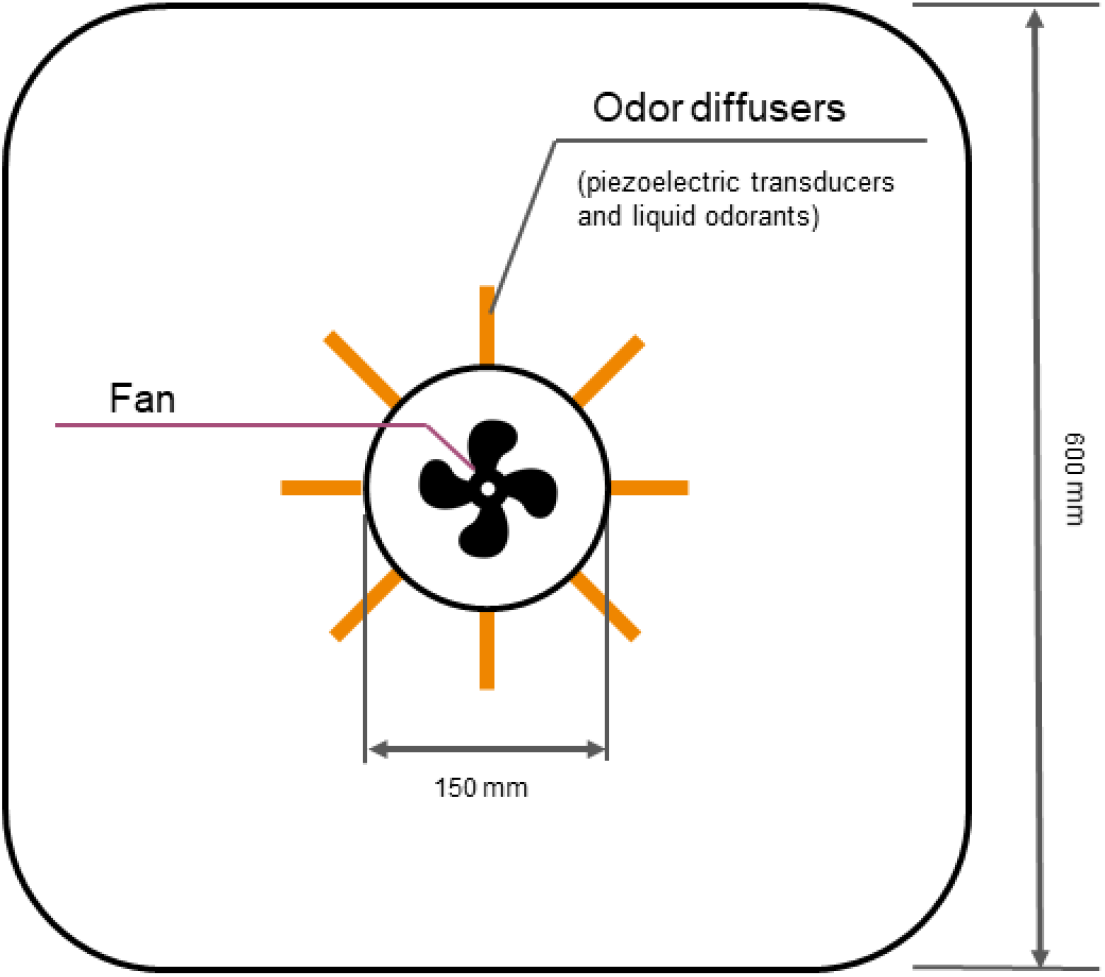
The schematics of olfactory display operation. The airflow was driven by a fan whereas liquid odorants were released under the control of piezoelectric transducers.

The experiments were conducted in a cubic enclosure (2.5 x 2.5 x 2.5 m), where a constant stream of air descended from the ceiling (Fig. 2). The participant was seated in an armchair in the middle of the room. A projector displayed visual stimuli on a screen mounted in front of the participant (Fig. 3). The olfactory display was mounted above the participant (Fig. 4 and Fig 5). The airflow for the olfactory display was 187 m^3^/hr For the outcoming air, the airflow was 580 m^3^/hr. The airstream was constant throughout the experiment. During the intervals between odorant delivery, the fresh air was delivered into the room. A video showing odor delivery process is available in the supplementary materials.

**Figure 2.**
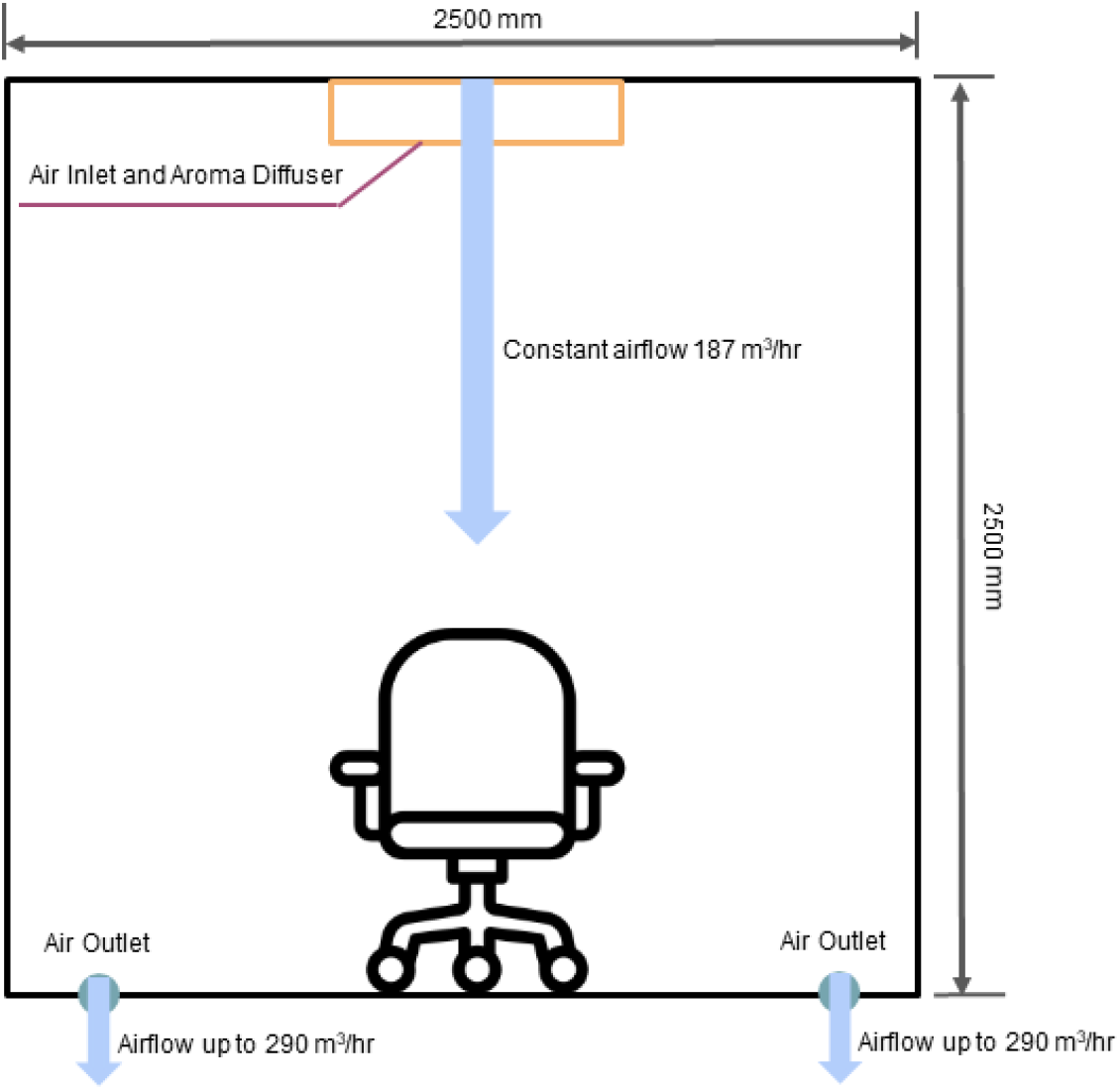
Schematics of the experimental setup. The participant’s chair was placed in a cubic enclosure, Odors were delivered from the ceiling, and the exit points for the airflow were located at the floor.

**Figure 3.**
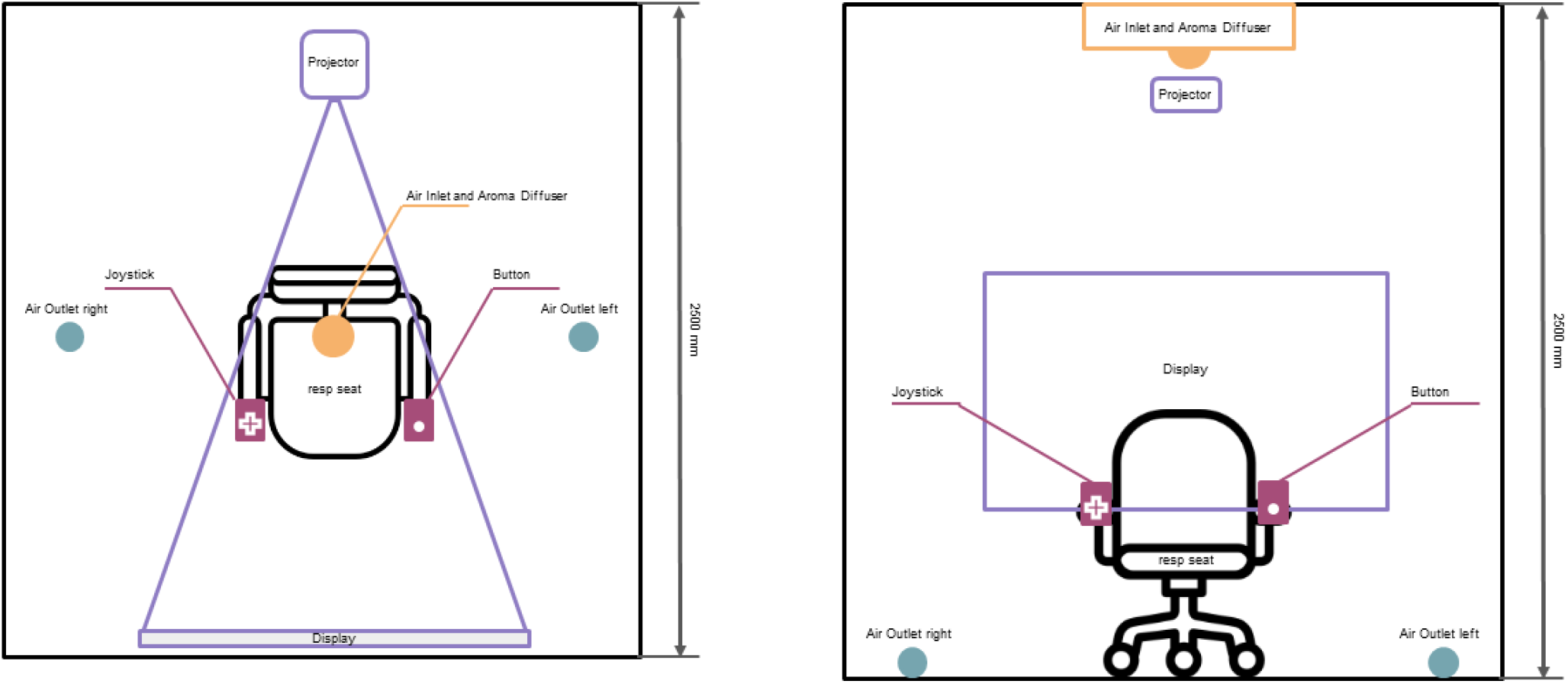
Schematics of the experiment setup. The view from the top is presented on the left and the view from the back is presented on the right.

**Figure 4.**
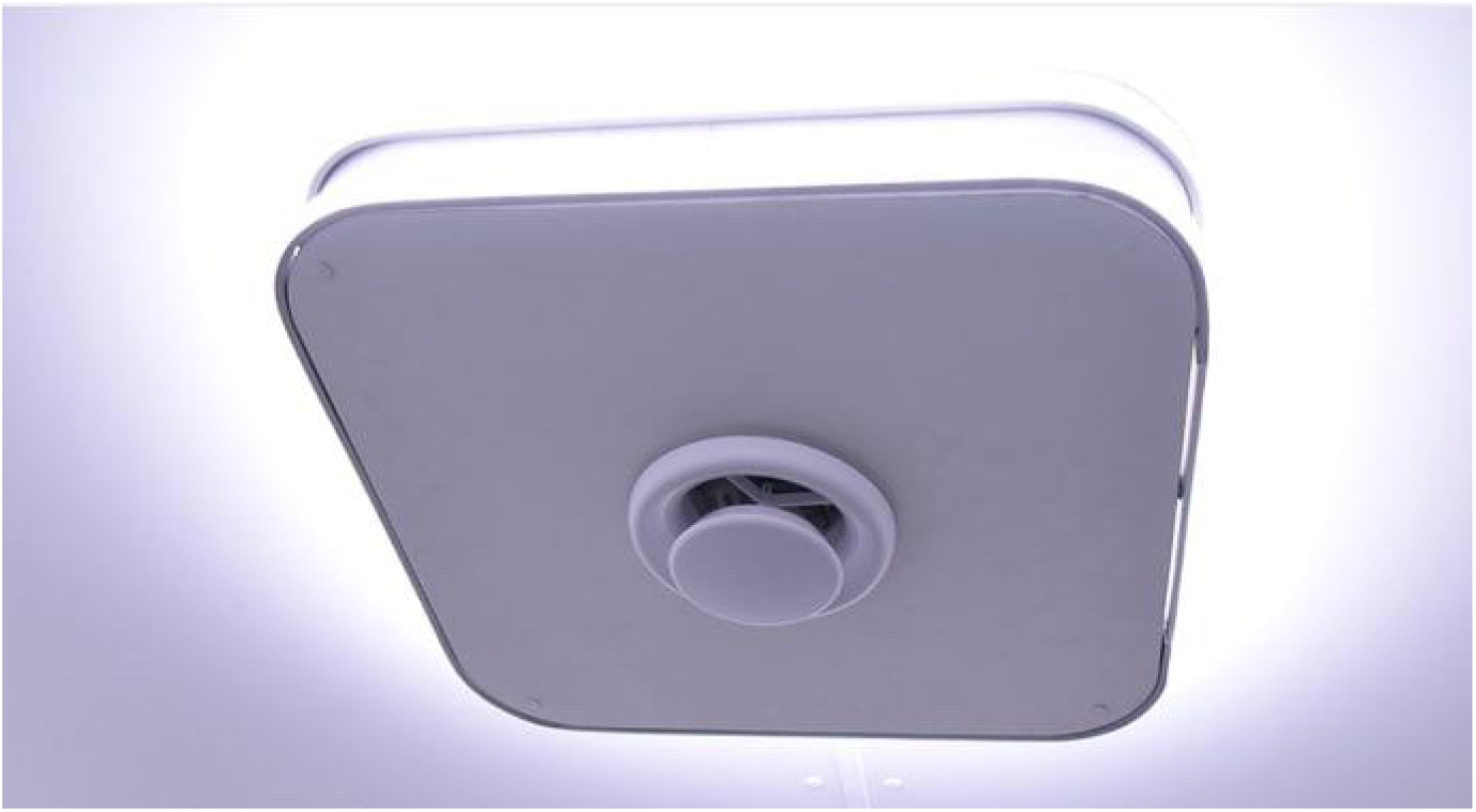
The olfactory display mounted to the ceiling.

**Figure 5.**
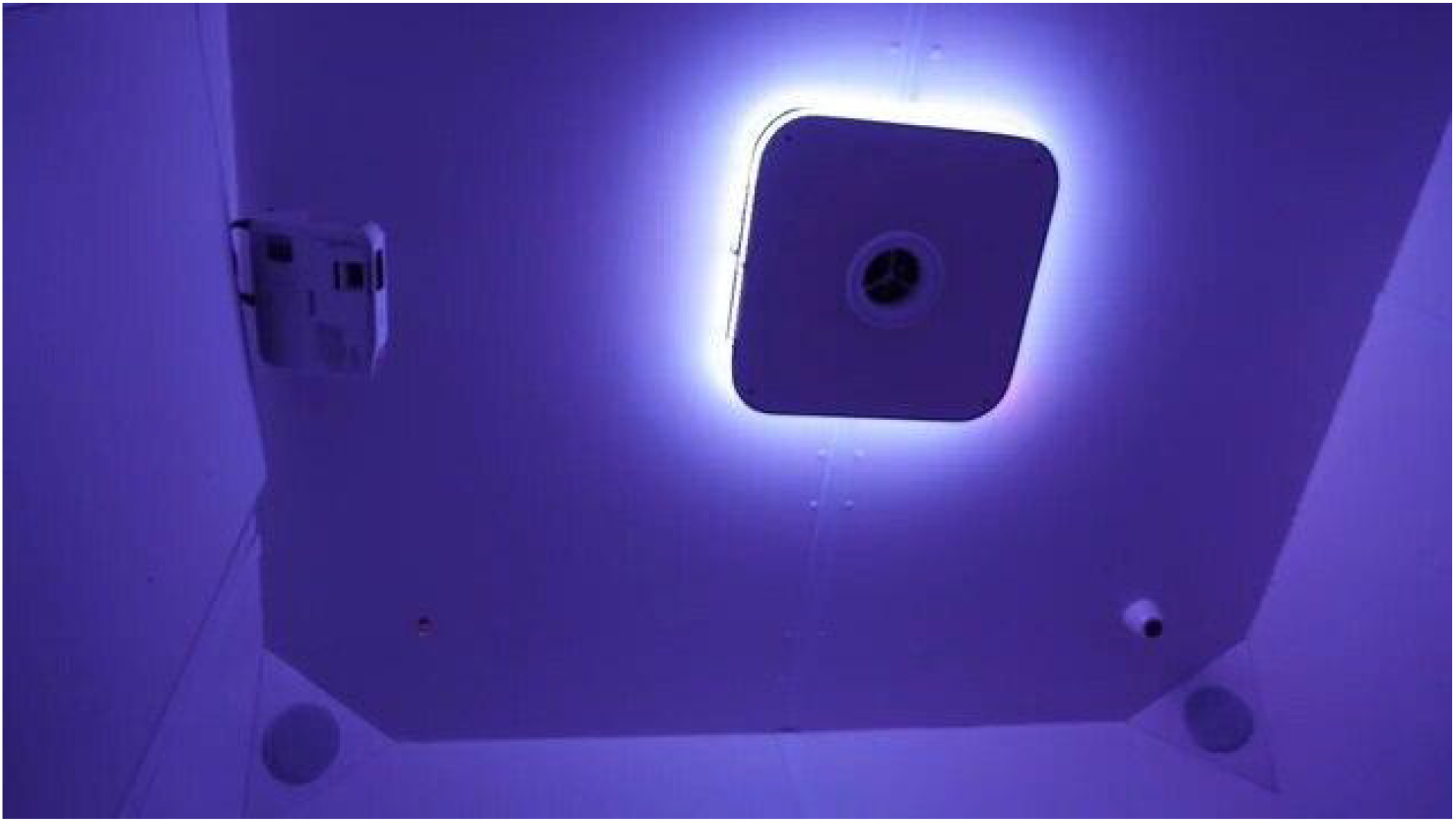
A photograph of the olfactory display depicting the room illumination coming from the ceiling.

### 2.2 Olfactory stimuli

The odor stimuli were dissolved in order to achieve viscosity necessary for stable evaporation with piezoelectric transducers. Ethanol-based solvent was applied to natural coffee, perfume vanilla and citrus essential oil. Four liquid stimuli were used: vanilla, coffee, citrus and odorless water (control stimuli). These flavors were chosen in a pilot study where we selected the odors that were familiar to the general public. In addition to the selected odors, we considered rose water and mint, but rejected these odors because rose water was unfamiliar to the Russian participants and they associated mint with taste rather than smell.

### 2.3 EEG and respiration registration

EEG data were collected with a Smart BCI system. 19 electrodes were positioned on the scalp according to the International 10–20 system with A1+A2 ears reference. Respiration data were collected with TRSens temperature sensor for nasal-oral breathing and KARDi2-NP polygraph amplifier. Data sampling was synced with the event markers for the button presses.

### 2.4 Joystick setup

Participants operated a two-dimensional joystick to report the perceived odors. We adjusted an arduino joystick for this purpose. The joystick was placed in a box that limited its movements to two dimensions.

**Figure 6.**
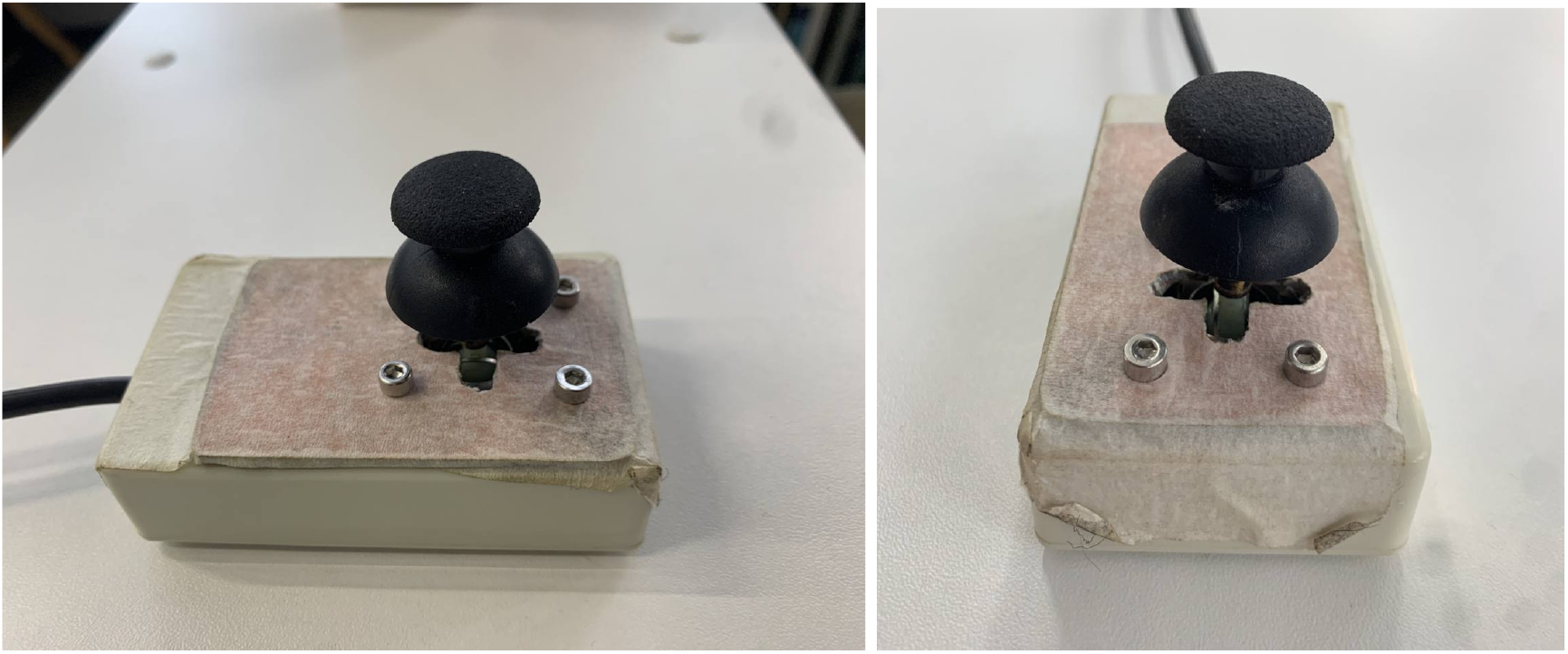
The two-dimensional joystick used for reporting the perceived smells. The joystick was a modified Arduino joystick which was in a frame that limited movements to four possible report directions.

The subjects initiated each trial by pressing a button with the left hand, which started the sequence of task events and synced the system components. A custom signal multiplier distributed the signals from the button to three devices: the computer that ran the experimental sequence and controlled the olfactory display, the EEG amplifier, and the respiration amplifier (Fig. 7).

**Figure 7.**
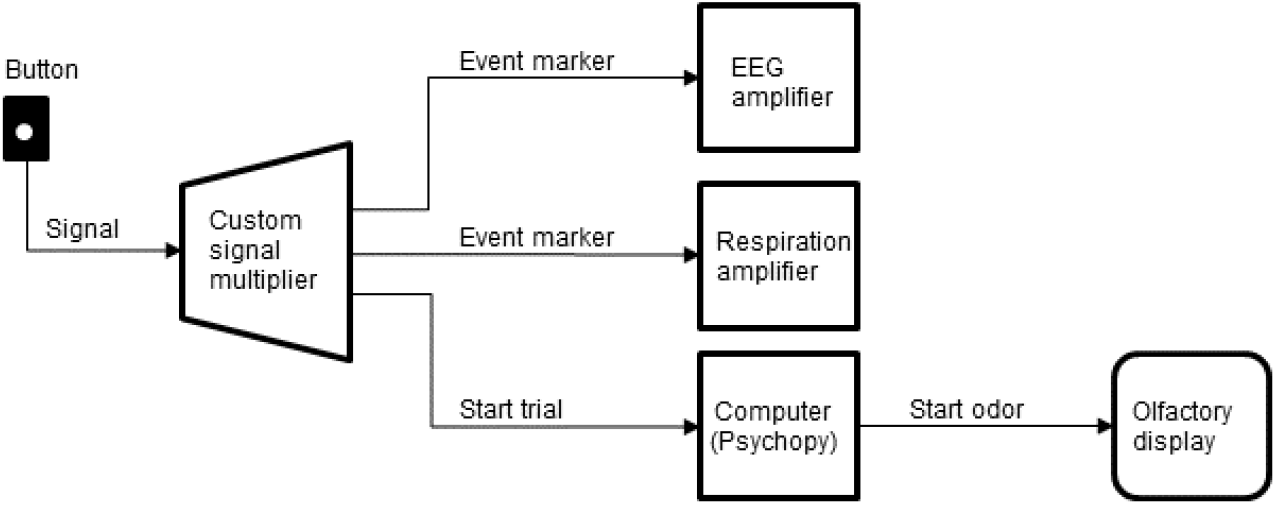
Schematics of the signal connection from the button to the other devices.

## 3 Methods

### 3.1 Subjects

All subjects signed informed consent forms prior to the experiment. The experiments were performed under the ethical approval from the Higher School of Economics Institutional Review Board (decision from 15th of February 2021). The participants did not receive any financial reward for this experiment.

We collected data in 17 healthy participants (age 22-44, median age = 31, females=7). One subject’s EEG and respiratory data were excluded because the system malfunctioned during the recordings. All subjects filled the forms before and after the experiment. The first form included general sociodemographic information and some specific questions regarding olfaction. Four participants stated that they have a olfaction-related hobby (perfumery, winery etc). None were professionally employed in fields that require special olfactory skills. None of the participants reported anosmia. One participant reported a broken nasal bone, which did not alter her odor perception.

### 3.2 Experiment task

The task was an instructed-delay task that required subjects to smell an odor and then transform this perception into a pointing movement performed with a joystick. By the experiment design, each odor (including no-smell condition) was associated with a visual label: a square, circle, triangle, or a star (Fig.8). Accordingly, the subjects reported their smell assessments by pointing to the appropriate label. Odor-label pairs were randomly generated for each participant and remained constant during each experimental session.

**Figure 8.**
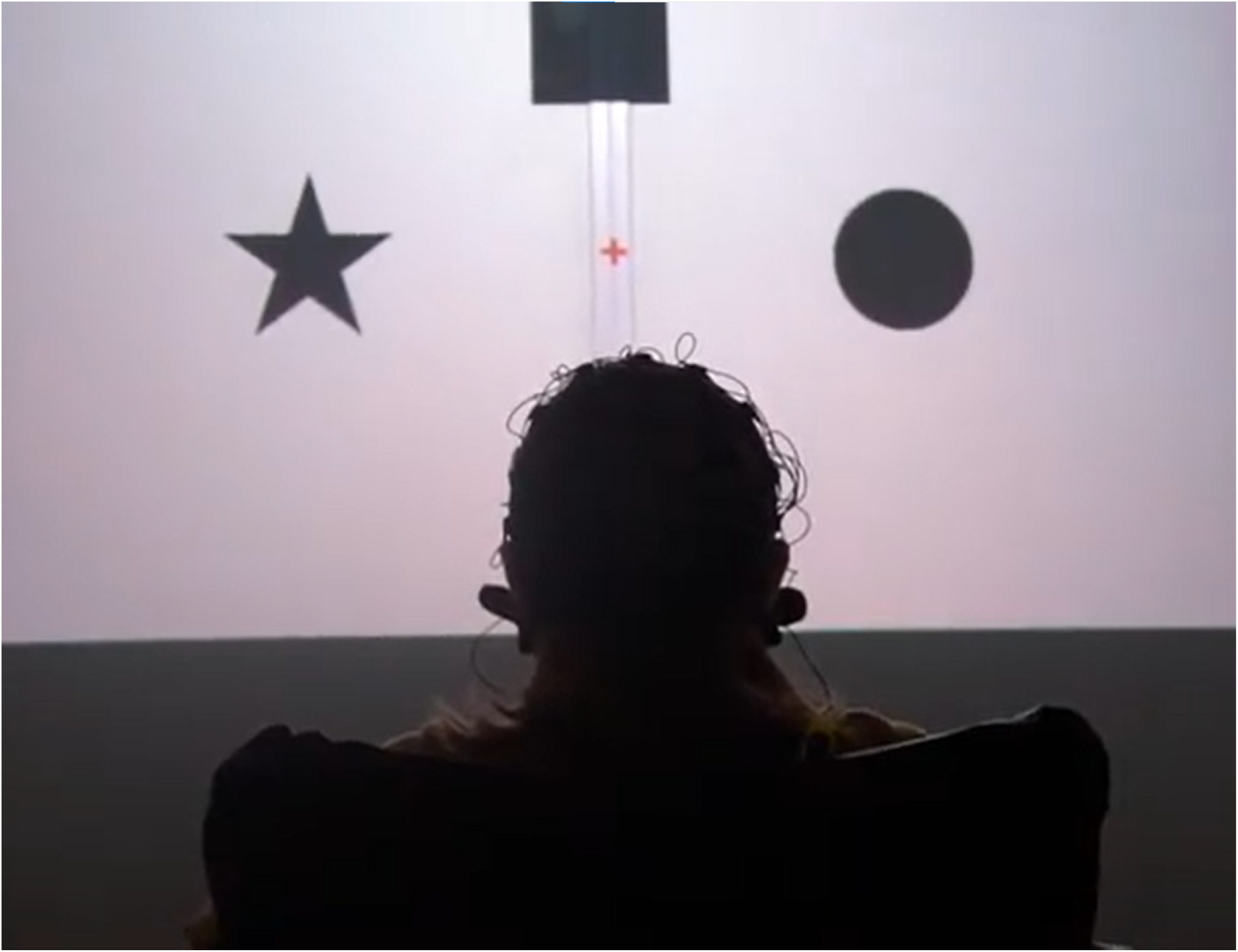
A photograph showing the participant sitting in front of the screen. The subject is viewing the stimuli associated with particular odors.

Following the training (40 trials, 10 x odor, random order), an odor discrimination session was run (80 trials, 20 x odor, random order). After starting a trial, participants were instructed to hold their breath until the fixation cross turned green. Next, they pressed a button. Odor delivery started immediately after the button was pressed. Following a 2-s delay, the fixation cross changed color from red to green and the participant made the first inhale. After the subject assessed the odor for the subsequent 10s, four objects appeared on the screen (at 0, 90, 180, and 270° positions); one of them represented the correct response. With this design, EEG and respiratory data were collected throughout all four task epochs: (1) no odor, (2) odor discrimination without any motor preparation, (3) motor preparation, and (4) peri-movement interval. After the entire experimental session was completed, participants verbally described their impressions of the odors delivered and other aspects of the task (Fig. 9).

Participants were instructed to avoid unnecessary movements during the trial and use time before the next trial if there was a need to flex the neck or hands or conduct any other movement. These settings minimized the artifacts in the EEG recordings. The duration of the experiment varied across participants because they were allowed to make short breaks between the sessions.

**Figure 9.**
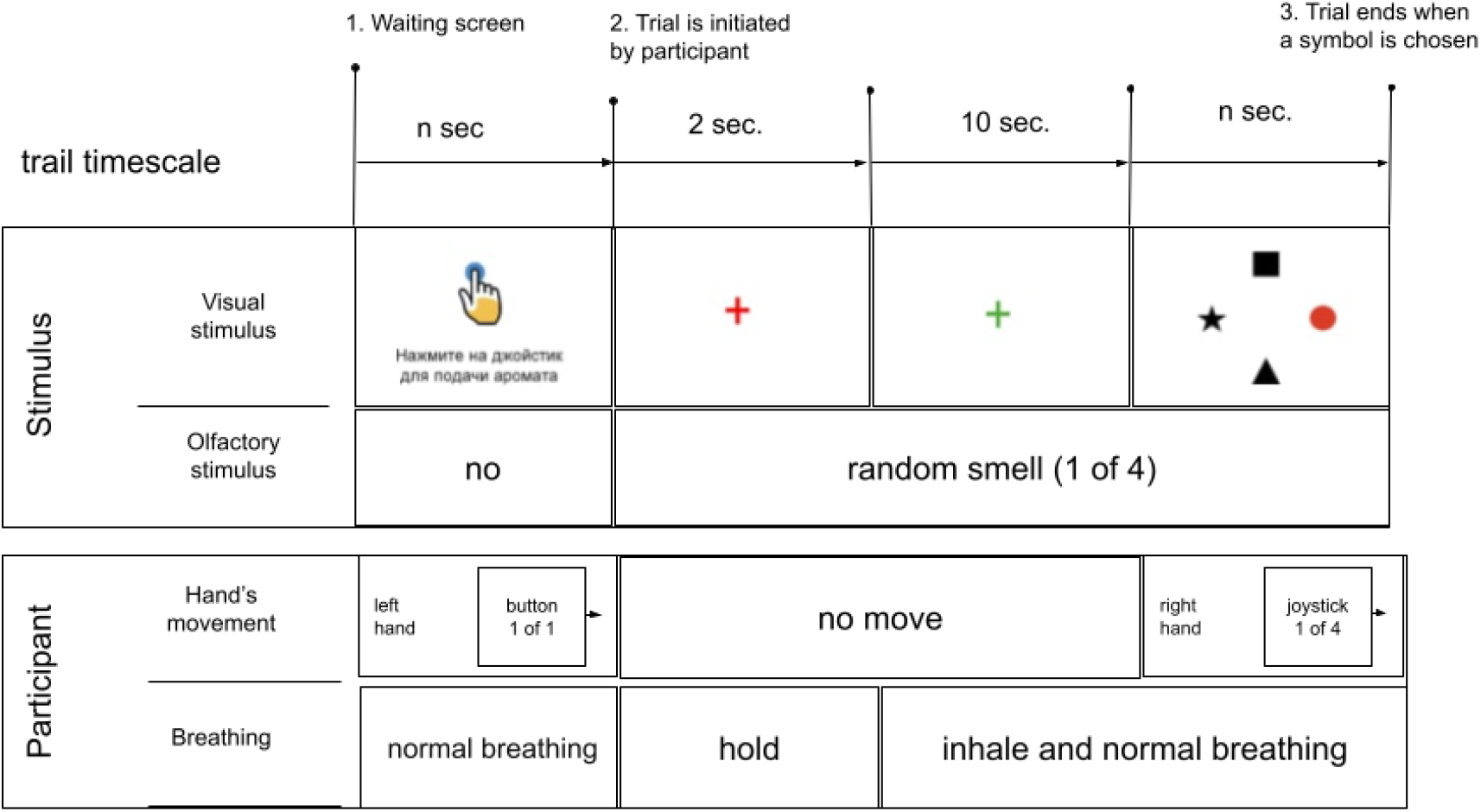
Schematics of the experimental sequence. In the beginning of each trial, an instruction (written in Russian) was shown on the screen: “Press the joystick for aroma delivery”. Next, the subject pressed the button to start the sequence of task events. A red cross appeared on the screen for 2s; the subject visually fixated the cross and held their breath. An odorant was delivered during this period. The cross then changed color into green, which instructed the subject to start inhaling. The subject continued breathing with a comfortable pace for 10s. Finally, a display with four items was shown on the screen, and the subject pointed with the joystick to the item that corresponded to the perceived odor. If the session was a training session, the correct item was highlighted with red color. During the discrimination that followed all items were colored black.

### 3.3 Respiratory data preprocessing

All participants were instructed to start inhaling when the fixation cross turned green. An algorithm was developed to detect the exact moment the subject started to inhale. The algorithm utilized a sliding window (1 second width), and the inhalation start was determined as the curve deviation from a stationary value. Some participants failed to hold their breath on some trials and/or the inhalation onset was not sufficiently abrupt to be detected by the algorithm. Thus, in an individual with a broken nasal bone, only 58 inhalation onsets could be detected out of 120 algorithmically or by visual inspection.The oldest female participant (39 years) had a breathing pattern where we could not detect many of the inhalation onsets. The algorithm was adjusted to avoid false-positive detections. The number and percentage of detected inhales for each subject are given in Table 1.The median number of detected inhales was 106.5 out of 120 total trials.

**Table 1.** Detection of inhalation onsets in different subjects.

| **Subject** | **# detected inhales**<br>**(out of 120)** | **% detected inhales** |
| --- | --- | --- |
| A4SHD_21F | 120 | 100.0% |
| V1XS8_31M | 119 | 99.2% |
| 9V6NM_36M | 118 | 98.3% |
| ZU9B3_34M | 118 | 98.3% |
| ZNH6Y_22M | 118 | 98.3% |
| 5V005_22M | 114 | 95.0% |
| 6K0UR_33M | 111 | 92.5% |
| 24WAG_31M | 111 | 92.5% |
| FK7CK_40M | 102 | 85.0% |
| VRWV7_31F | 100 | 83.3% |
| 7PF8Z_36F | 89 | 74.2% |
| O9ZEA_23F | 88 | 73.3% |
| VFDSF_20M | 85 | 70.8% |
| 3KACQ_44M | 76 | 63.3% |
| 99CEM_22F | 58 | 48.3% |
| 1R912_39F | 39 | 32.5% |

### 3.4 EEG preprocessing

Independent component analysis was used to remove EEG artifacts. The algorithm generated the components based on a mutual information analysis. The artifact removal was performed under visual inspection. A low-pass filter with a 45-Hz cutoff, and a high-pass filter with a 2-Hz were used.

## 4 Results

### 4.1 Behavioral results

Each subject performed 120 trials, with 40 training trials and 80 discrimination trials. The majority of participants (12 out of 17) successfully reported the presented odor in more than 90% of the trials. The median accuracy was 93.8% with standard deviation of 14.5%. The weakest result (35 correct reports out of 80 trials, 32.9%) was in the oldest male participant (age = 40). His performance accuracy was still significantly above the chance level of 25%. Performance accuracy across the participants is shown in Table 2.

**Table 2.**
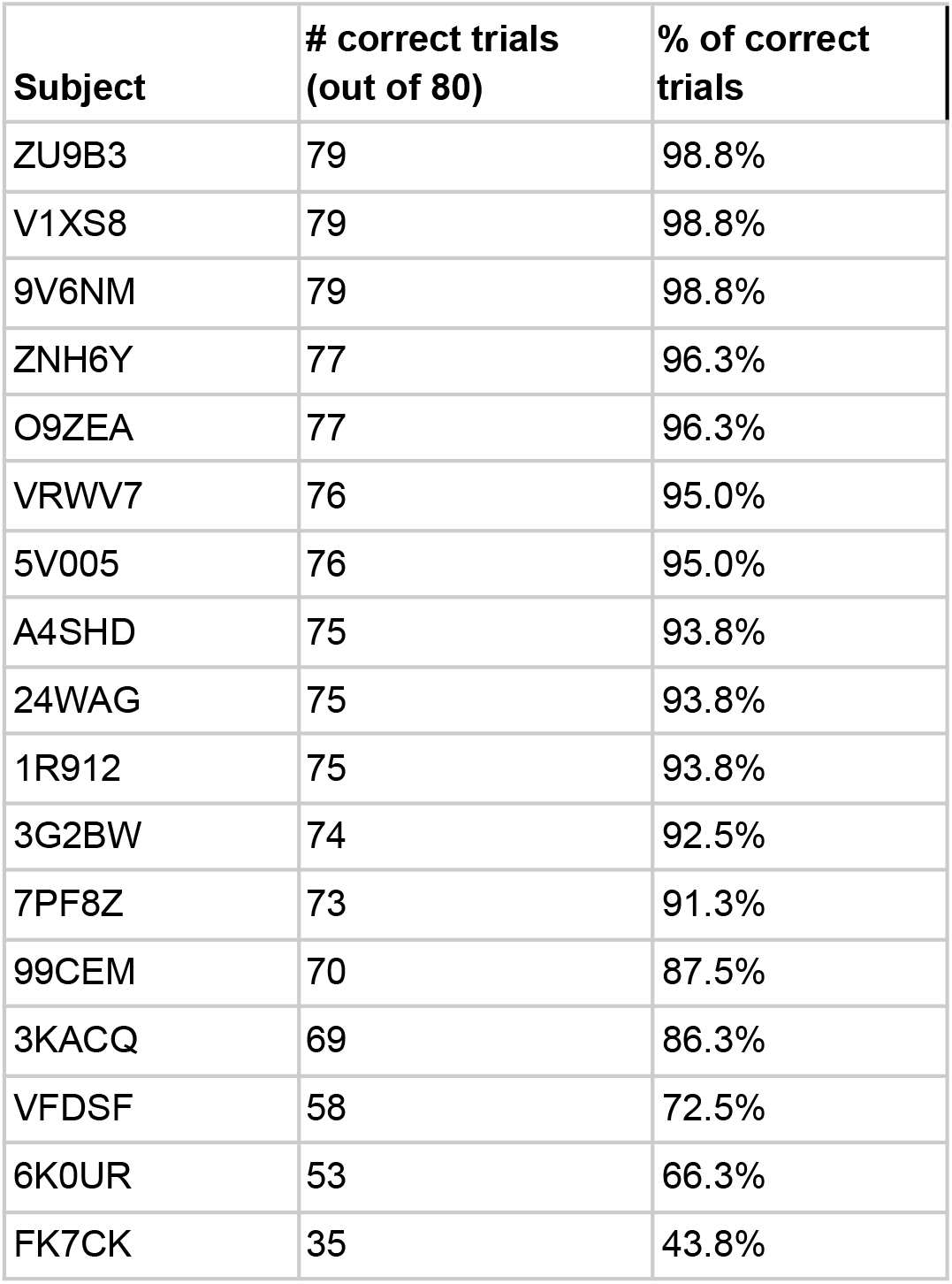
Performance accuracy across the participants.

| Subject | # correct trials<br>(out of 80) | % of correct<br>trials |
| --- | --- | --- |
| ZU9B3 | 79 | 98.8% |
| V1XS8 | 79 | 98.8% |
| 9V6NM | 79 | 98.8% |
| ZNH6Y | 77 | 96.3% |
| O9ZEA | 77 | 96.3% |
| VRWV7 | 76 | 95.0% |
| 5V005 | 76 | 95.0% |
| A4SHD | 75 | 93.8% |
| 24WAG | 75 | 93.8% |
| 1R912 | 75 | 93.8% |
| 3G2BW | 74 | 92.5% |
| 7PF8Z | 73 | 91.3% |
| 99CEM | 70 | 87.5% |
| 3KACQ | 69 | 86.3% |
| VFDSF | 58 | 72.5% |
| 6K0UR | 53 | 66.3% |
| FK7CK | 35 | 43.8% |

To assess the discrimination performance for different odors, a confusion matrix was calculated, where the matrix rows represented odors being presented, the columns represented odors being reported, and the values were report counts (Fig. 10). In this analysis, each subject performed 20 discrimination trials for each odor, so there was a total of 340 trials per odor for all 17 subjects. The confusion matrix showed an overall good performance for all subjects. The mean success rate was 93.8% with standard deviation of 14.5%. The best discrimination performance was for vanilla odor (312 correct responses out of 340 trials) and errors occurred most often for citrus (52 errors out of 340 trials).

The statistics of odor discrimination was assessed with a chi-squared test applied to the confusion matrix (Table. 3). Additionally, pre-whitened Mann-Kendall test for all the correct responses from each subject per each trial (Fig. 11) revealed the absence of a significant trend (z=-1.5609, p=0.1186), as well as the test of the mean response time (z=-0.3302, p=0.7412, Fig.12). The statistical testing of individual response times demonstrated significant decrease in response times for some of the participants (Fig. 12): 1R912 (z=-3.9793, p<0.001), 5V005(z=-3.1157, p<0.05), 9V6NM (z=-3.6661, p<0.001), FK7CK (z=-4.7583, p<0.001). The only significant increase was found for participant 7PF8Z (z=2.1251, p<0.05).

**Figure 10.**
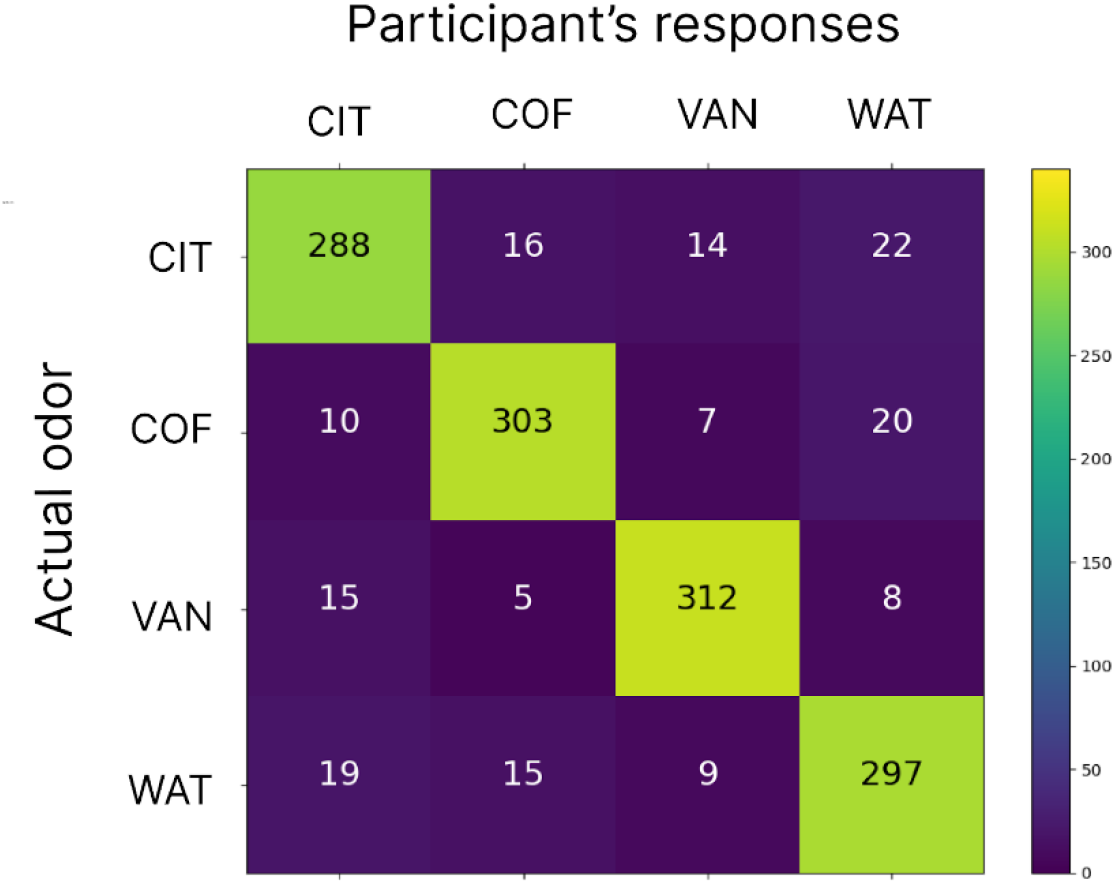
The confusion matrix for participant’s responses and actual odors being delivered. Numbers indicate the counts of responses for different delivered odors. Each odor was delivered 340 times (20 trials per odor for 17 subjects). The color scale is from blue (minimum) to yellow (maximum). CIT, COF, VAN and WAT are citrus, coffee, vanilla and water, respectively.

**Table 3.** The results of the chi-squared test for the confusion matrix demonstrating the independent perception of different odors.

| Stimulus | Power divergence | p-value |
| --- | --- | --- |
| citrus | 675.590 | <0.001 |
| coffee | 750.263 | <0.001 |
| vanilla | 800.339 | <0.001 |
| water | 680.746 | <0.001 |

Following the discrimination session, subjects were asked to fill in the form and provide names for the odors that were delivered. All participants correctly named odorless water and the majority of them (10 out of 17) named coffee. Yet only 5 and 3 subjects could correctly name citrus and vanilla odors, respectively. All subjects except for one reported that their olfactory discrimination abilities did not change throughout the experiment. One subject, the oldest male in the group, made more mistakes in the second half of the experiment.

**Figure 11.**
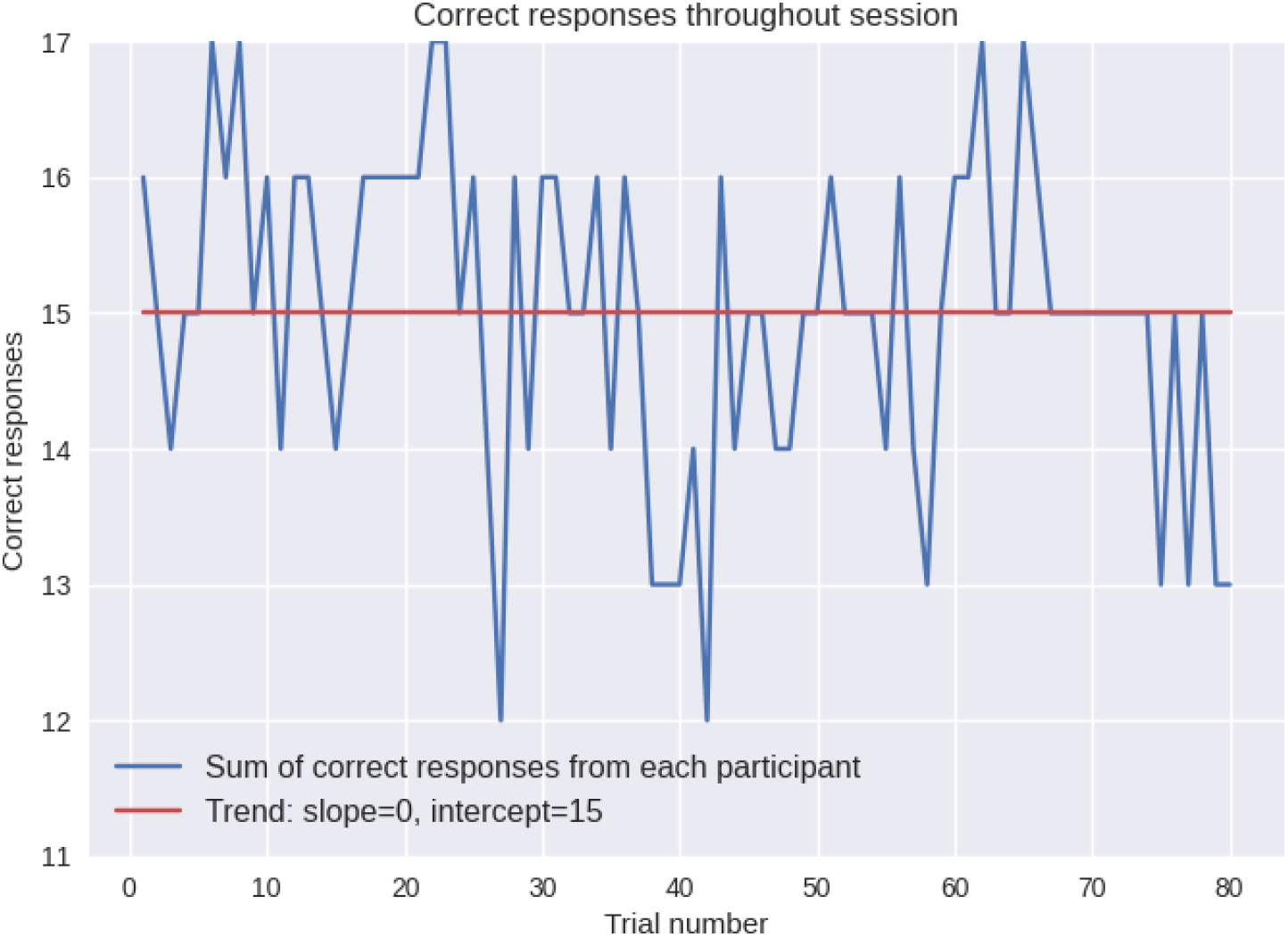
Number of subjects who responded correctly for each trial. No significant trend has been revealed. There was no significant change in the accuracy during the experiment.

**Figure 12.**
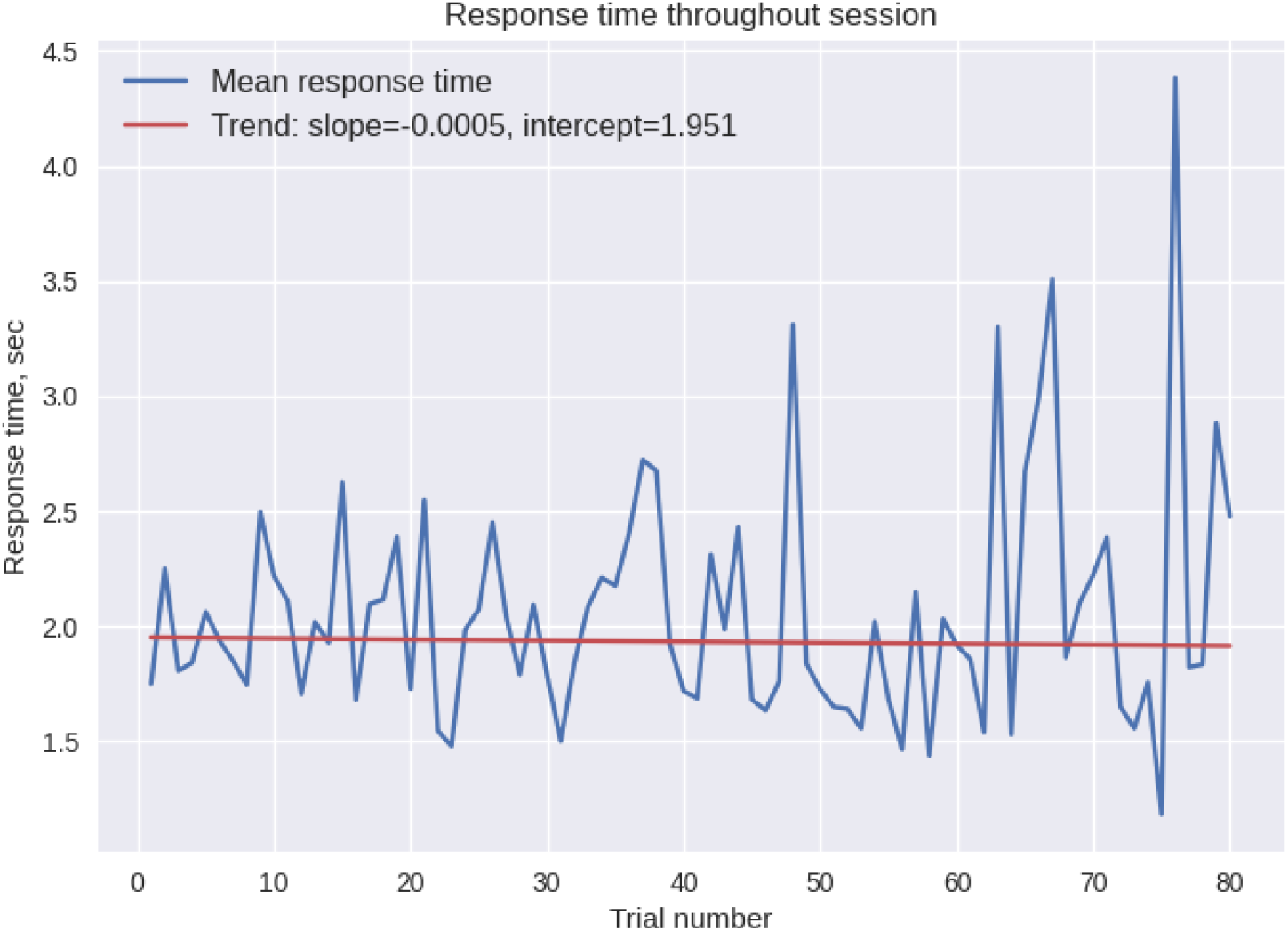
Mean response time across all participants for each trial. No significant trend has been revealed. There was no significant change in the response time during the experiment.

**Fig. 13.**
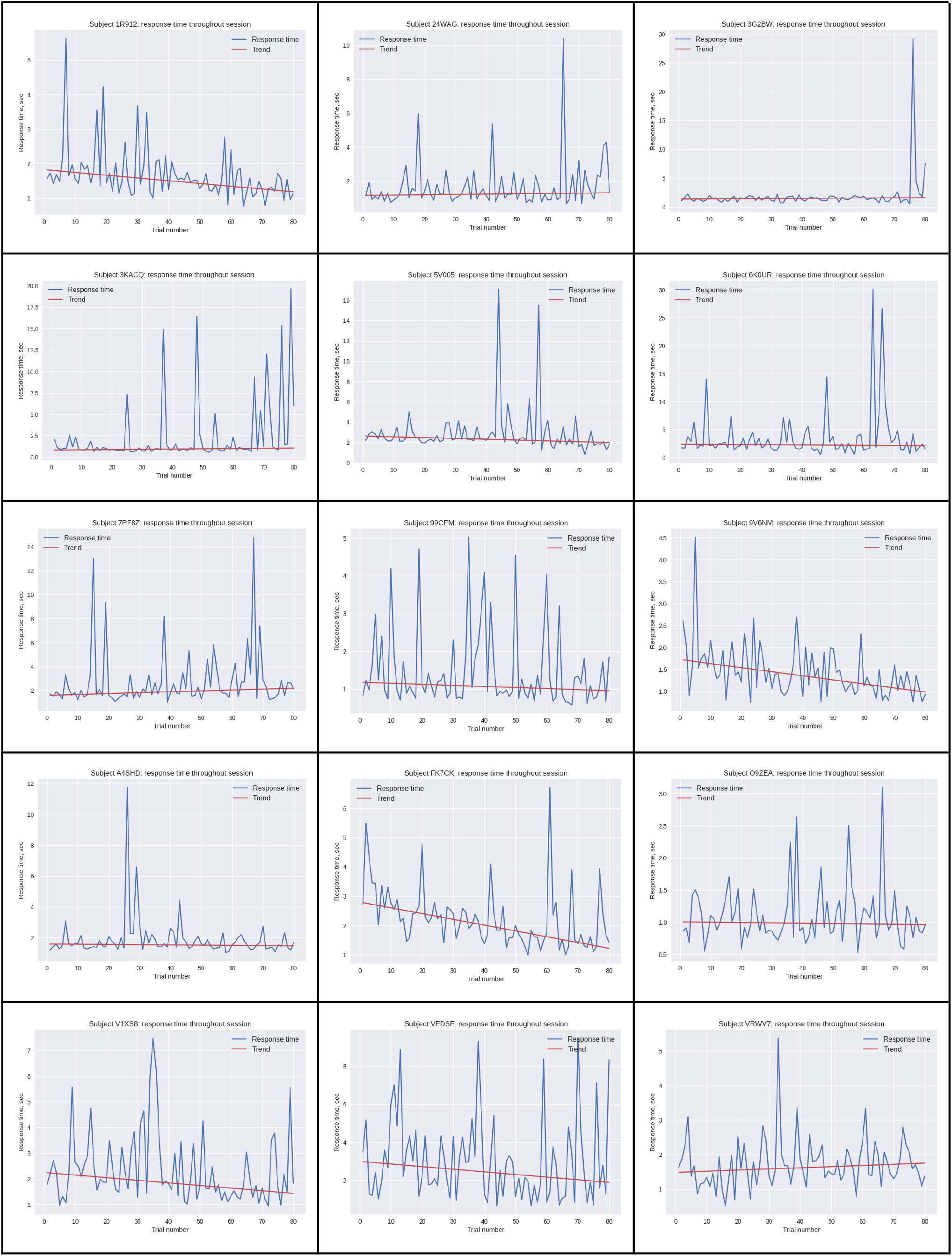
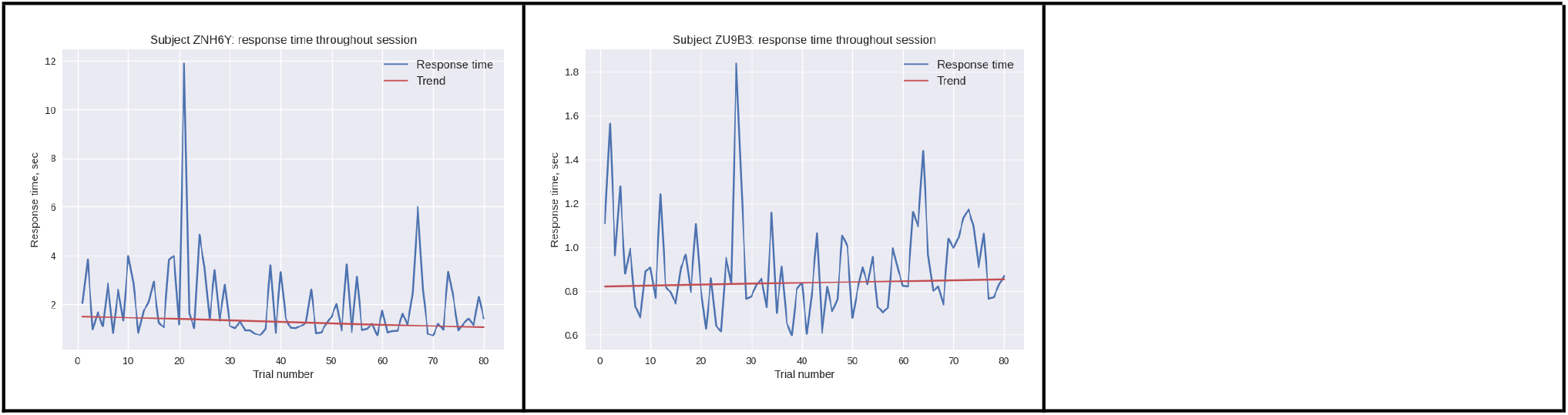
Individual response times as a function of trial number.

### 4.2 Olfactory related components

In the analysis of EEG patterns representing odors, we focused on determining an independent olfactory-related component. We examined the 7-s epochs following the inhalation onset (within a 10-s period of odor processing and prior to symbols being displayed on the screen, thus prior to any motor activity). When an ICA was applied to low-pass (below 20Hz) filtered EEG, we observed similar odor-related components in six subjects. The activity of these components was localized around the C4 electrode (C3 for one subject) with peak activity at the alpha-band frequency (10-12Hz). Figure 14 demonstrates localization and spectrum analysis of these components.

Responses to odors were detected in the recorded EEG activity. By task design, perception of olfactory stimuli started with inhalation onset, that is the EEG evoked responses correspond to both olfactory discrimination and the act of inhalation. If the evoked response reflected only motor activity there should have been no difference in evoked responses for different odors. Pairwise comparison of this component of EEG activity for processing of different odors demonstrates significant differences in the components’ spectral power for some odor pairs (p < 0.05). This was observed for the odors and odorless control stimuli, as demonstrated for participants O9ZE and V1XS8 in Figure 15 and Figure 16 accordingly. In some participants, significant differences were observed for spectral properties corresponding to perception of different odors, as demonstrated for participant A4SHD (Figure 17). Respiratory spectral properties demonstrated no difference for perception of vanilla and citrus. This difference in the EEG activity led us to conclude that this component represented not only motor elements of inhalation but also included the processes related to olfactory processing. These recordings were conducted prior to the joystick movement, so this activity cannot be related to hand movements.

**Figure 14.**
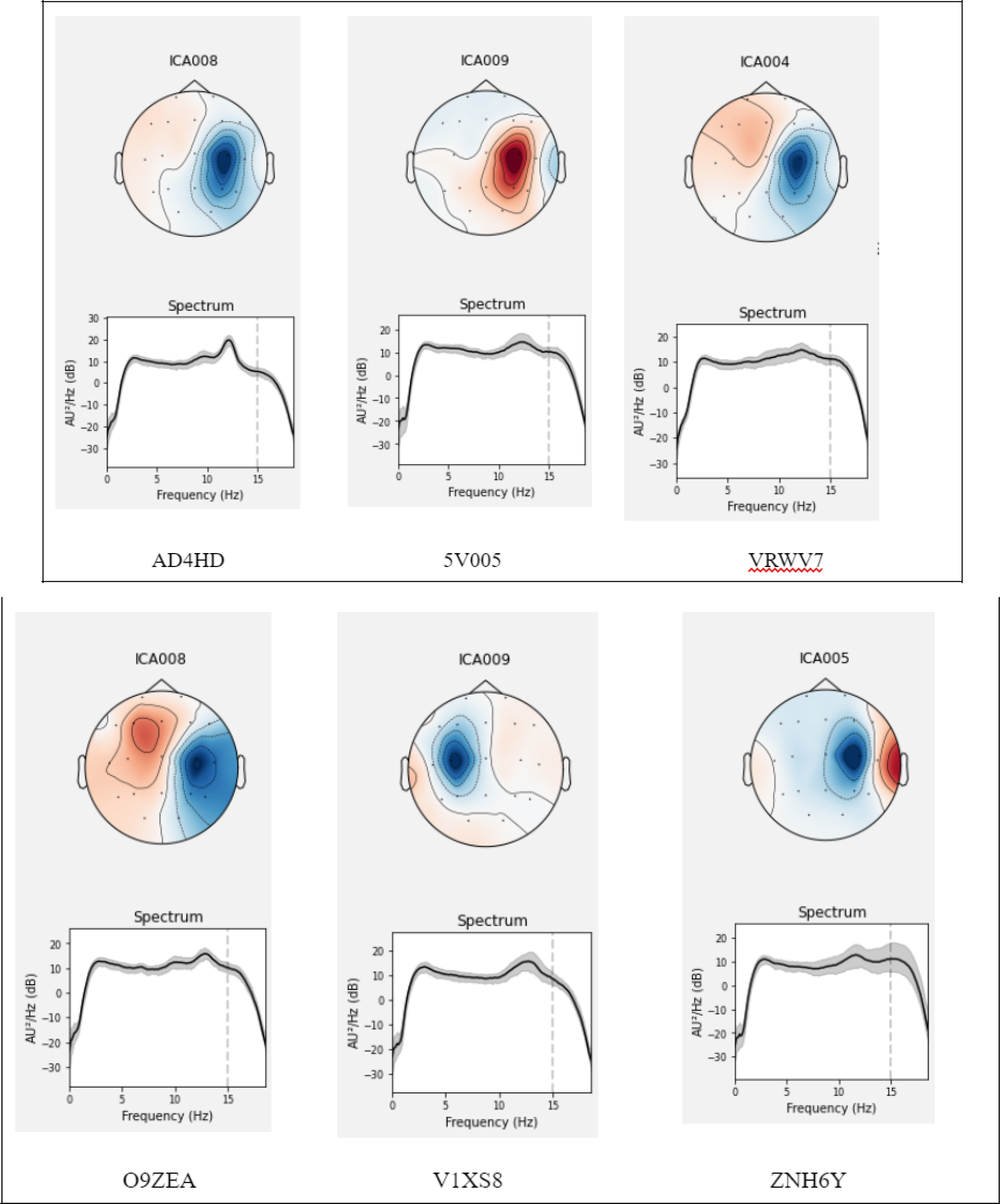
Location of the olfactory related component for six participants. The 7-s epochs after the inhale onset were analyzed, which preceded motor activity. ICA with the low-path filtering below 20Hz allowed to extract an olfactory related component localized around the C4 lead (C3 for one participant).

**Figure 15.**
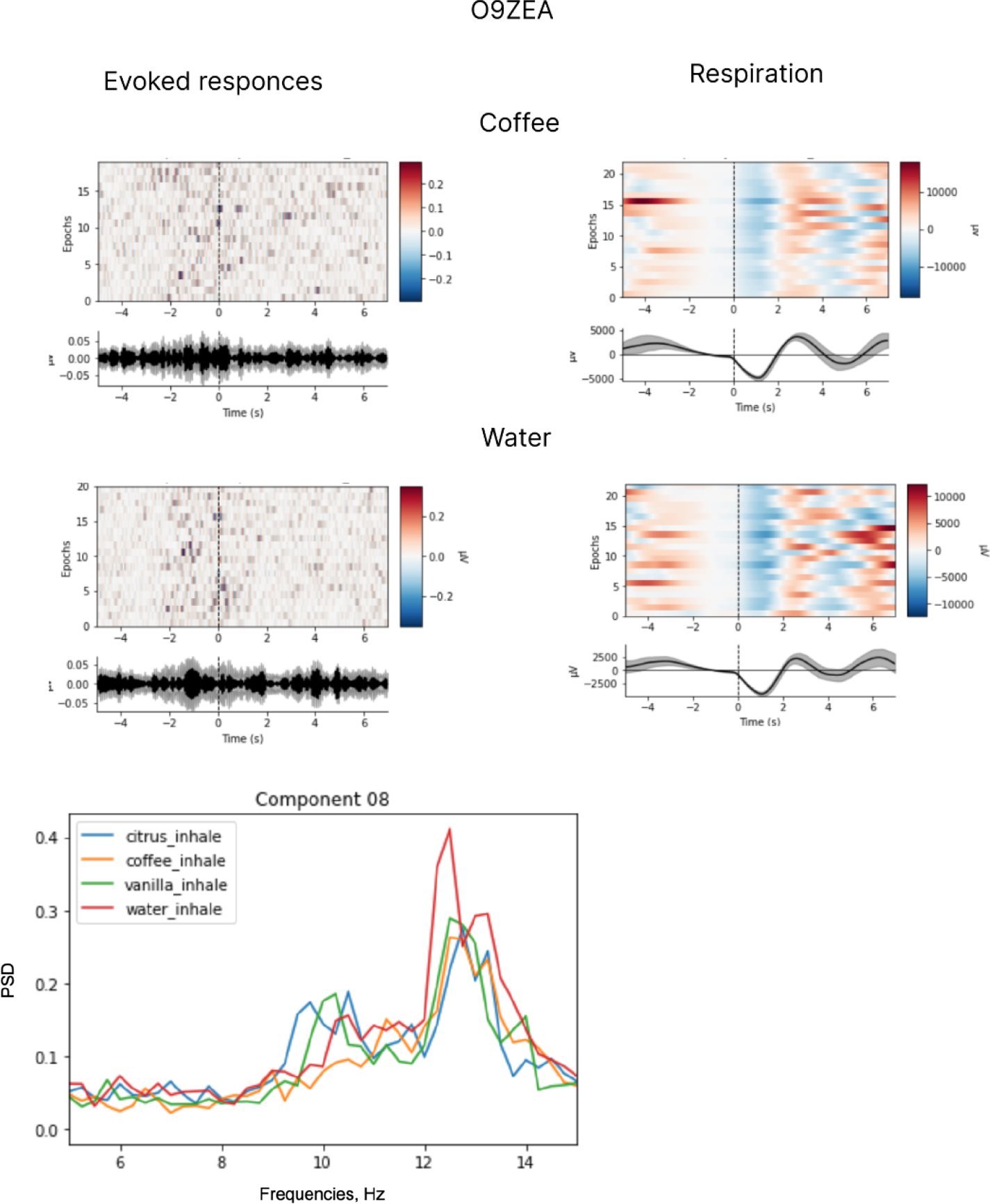
Evoked responses, respiratory data and spectral properties for coffee and water for participant O9ZEA. 7-ss epochs after the inhale onset were analyzed, which preceded hand movements. ICA with the low-path filtering below 20Hz (t=-2.2446, p<0.05 for peak around 10 Hz in ‘coffee’ and ‘vanilla’ conditions p value <0.05= 0.04 for t-test statistic applied at 10 Hz peak for conditions of coffee inhale and water inhale).

**Figure 16.**
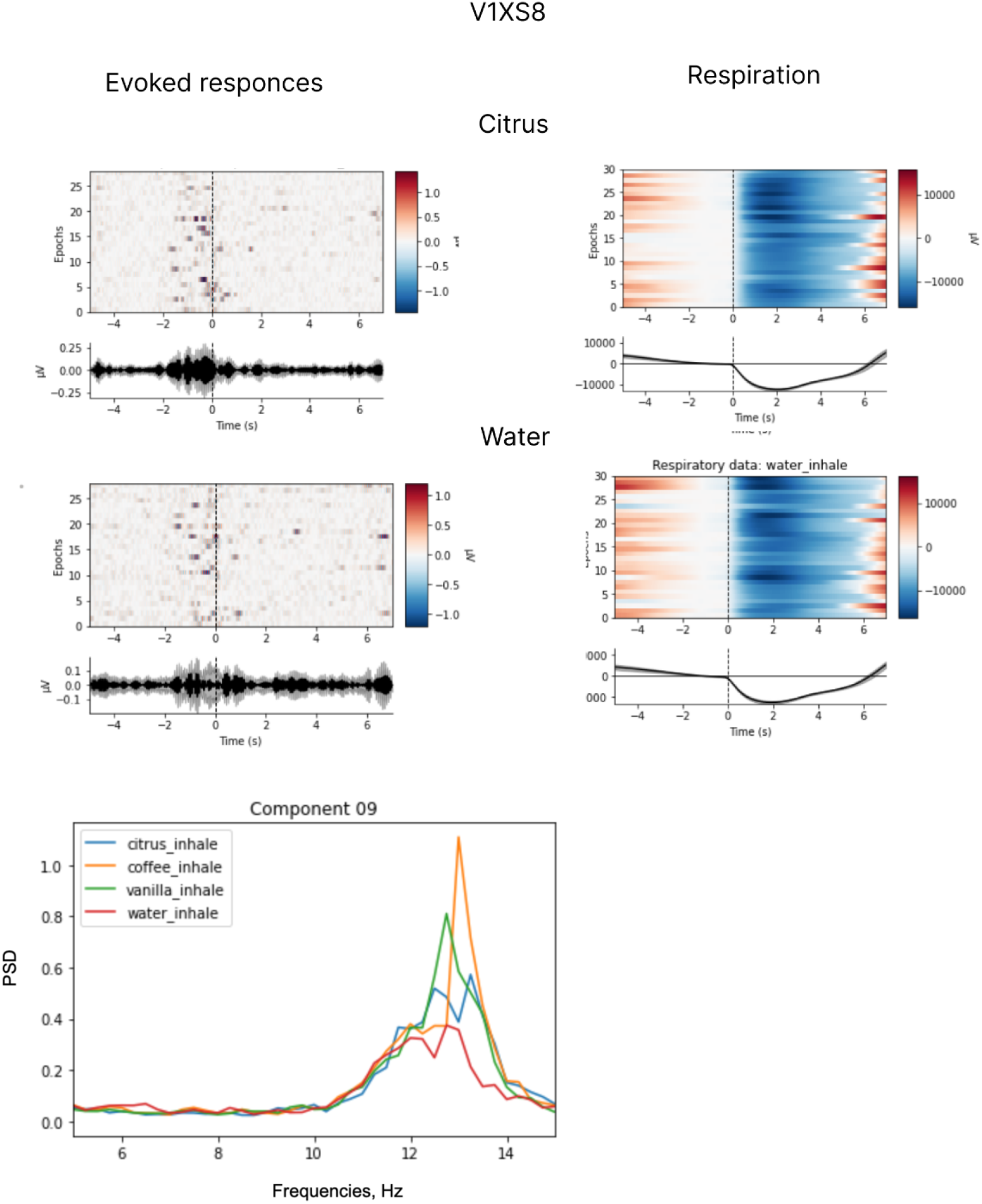
Evoked responses, respiratory data and spectral properties. Evoked responses and respiratory data for vanilla and citrus for participant V1XS8. The analysis was conducted for the 7-s epochs after the inhale onset, prior to hand movemnts. ICA with the low-path filtering below 20Hz (t=2.9174, p<0.01 for ‘citrus’ and ‘water’ conditions, t=2.232, p<0.05 for ‘vanilla’ and ‘water’ conditions).

**Figure 17.**
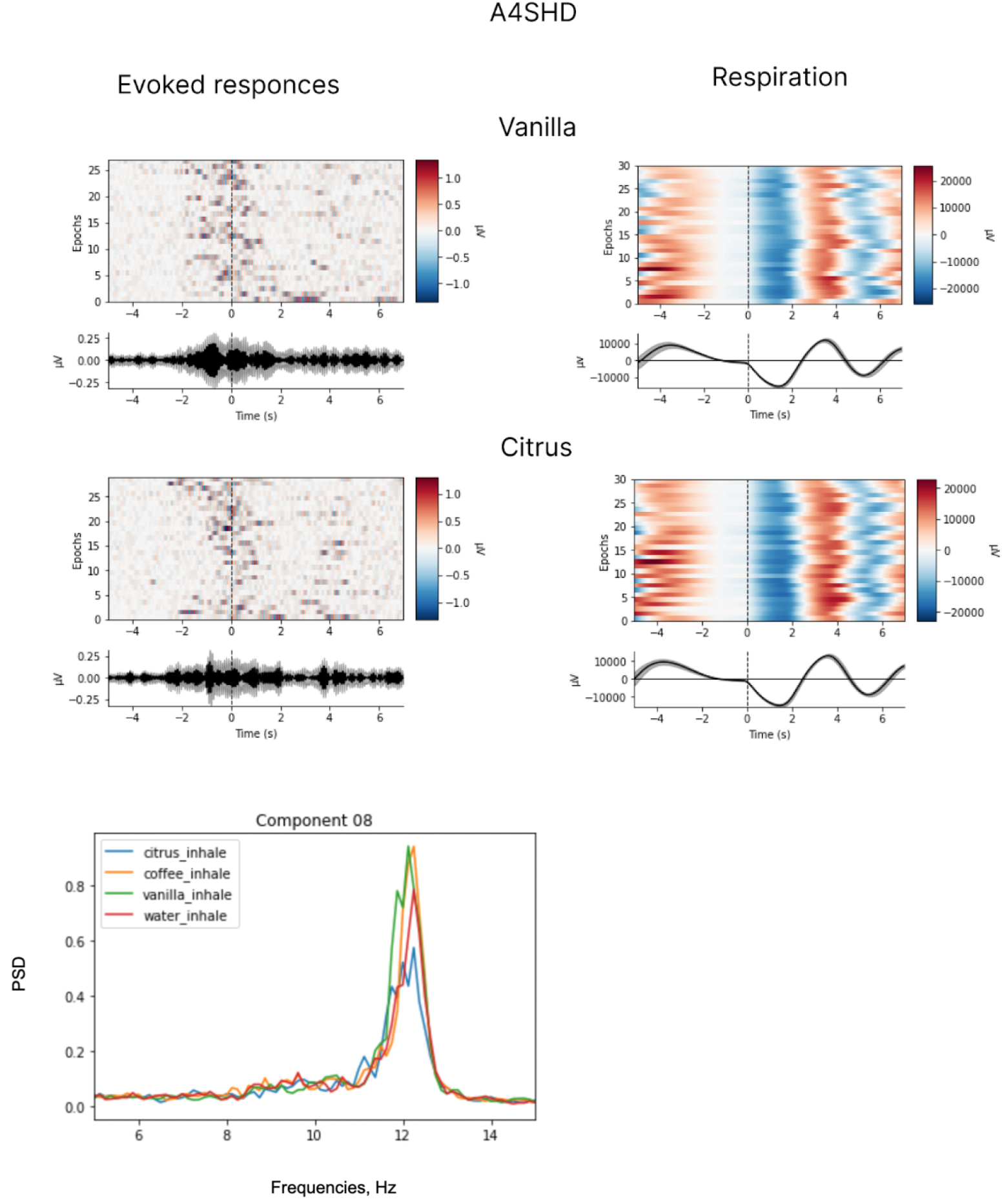
Evoked responses, respiratory data and spectral properties. Evoked responses and respiratory data for vanilla and citrus for participant A4SHD. Analyses were conducted for the 7-s epochs after the inhale onset, which preceded any hand movements. ICA with the low-path filtering below 20Hz (p value = 0.04 for t-test statistic applied at 10 Hz). (t=-2.2694, p<0.05 for ‘citrus’ and ‘vanilla’ conditions).

### 4.3 Olfactory related components in the instructed-delay task

Clear evoked responses were found for the final appearance of the set of response targets. Following the appearance of the targets, the subjects had to point to one of the labels with the joystick. As evident from the resulting grand average plot from the central channels C3, Cz and C4 (Fig.18), the P200 response was suppressed in cases when no odor was perceived (condition ‘water’).

**Figure 18.**
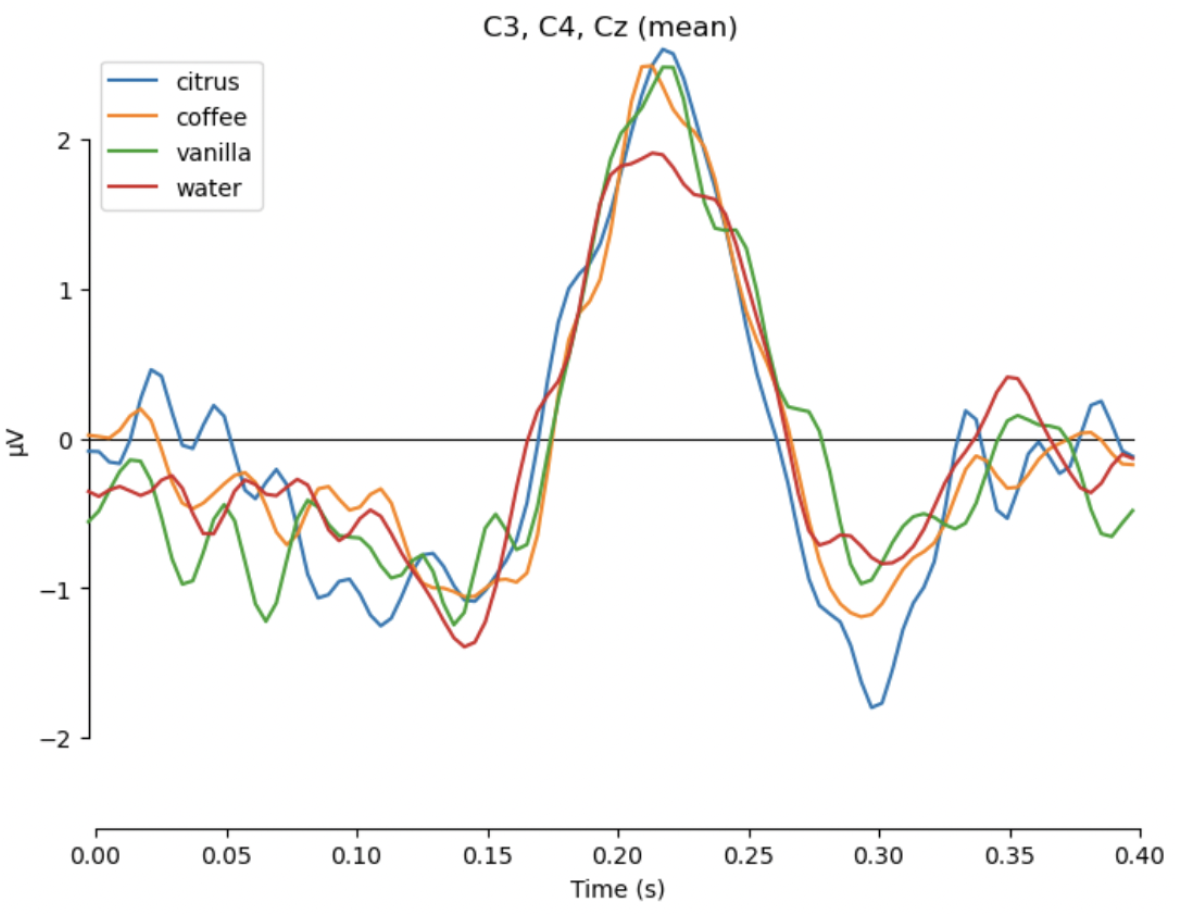
Evoked response to the presentation of final response targets, averaged across all subjects and channels C3, C4, Cz.

The pairwise t-test performed on the individual evoked responses and the mean amplitude around the peak of P200 demonstrated statistically significant differences between each of the odors and the control condition: t=2.78, p<0.01 for citrus; t=2.149, p<0.05 for coffee; t= 2.114, p<0.05 for vanilla.

Additional analysis of the peak-to-peak amplitude between P200 and N200 components revealed significant difference between ‘citrus’ and ‘water’ conditions: t=2.679, p<0.05, while the differences between ‘vanilla’ and ‘water’ were at the margin of statistical significance :t=1.754, p=0.086. The differences between ‘coffee’ and ‘water’ conditions were insignificant: t=1.338, p=0.18.

## 5 Discussion

We developed here a novel paradigm for the analysis of EEG patterns during olfactory processing in humans. The paradigm is an instructed-delay task where olfactory stimuli are presented using an automated odor display followed by a period of odor assessment and the final report made with a joystick. The odor display enabled controlled odor delivery. The settings resembled odor perception in the natural environment. In previous studies, odors were delivered manually (Aydemir 2017, Abbasi et al., 2019; Ezzatdoost, Hojjati and Aghajan, 2020; Zhang, Hou, Meng, 2019; Saha et al 2013) or using an olfactometers attached to the nostrils (Murphy et al, 2000; Rombaux et al, 2006; Invitto, Mazzatenta, 2019). In our settings, the olfactory display injected odors into the airstream. The stimuli were delivered in low concentration, close to the odors encountered natural environment.

The analysis of the behavior showed that human subjects performed the odor discrimination task well. Mean accuracy was 93.8%, which demonstrates that the proposed design is suitable for running olfactory-cognition tasks. The subjects continued discriminating odors for a long time, up to 1.5 hours (or 120 trials). This is somewhat surprising given the previous report that odor perception declines due to olfactory adaptation (Dalton, 2000). We observed adaptation in one subject, the oldest male in the group (40 yo). All other subjects did not demonstrate any signs of reduction of their ability to discriminate odors. We conclude that with a proper task human subjects are capable of perceiving and discriminating odors for a long period of time.

The ICA analysis of the 7-s epochs following inhalation onset allowed us to observe an olfactory related EEG component in six subjects. This component was typically located near the C4 electrode and peaked around 12 Hz. Since odor perception occurs during inhalation, this introduces a motor component related to the respiration. While there were no significant differences in the respiration pattern, pairwise comparison of this component of EEG activity for processing of different odors demonstrated significant differences for some-odor pairs (p value < 0.05). Such a difference in EEG activity led us to conclude that this component represented not only motor elements of inhalation but also included the processes related to olfactory perception. As such, this component should be further explored in order to develop a robust olfactory-based BCI system.

The P200 component, suppression of which was found in the control trials after the presentation of multiple-choice options, is known to be crucial for primary categorization and attention-related processes (Luck et al., 1994). The target stimuli elicited more pronounced P200. In the described setup, the control condition with no odor perceived could be considered as non-target, implying a different type of processing during the decision-making, namely reporting of stimulus presence as opposed to reporting its absence. Since the olfactory stimuli were presented until the instance of making the final decision, the higher amplitude of the P200 component could reflect an attentional access to the presented odor and its identification (which is absent in the case when the odor is not perceived). Thus, the P200 component can serve as a robust marker of odor identification initiated by the force-choice delayed task and can be used in olfactory-based BCIs.

## 8 Acknowledgments

Authors want to thank Grigory Gritsenko, Alexey Osadchiy and Ruslan Krashenkov for their helpful suggestions regarding study design, experimental design and data analysis.

Supplementary materials: video of the olfactory display and the air outlets https://youtu.be/-67AiXuXMxg

